# TTAPE-Me dye is not selective to cardiolipin and responds to main anionic phospholipids unspecifically

**DOI:** 10.1101/2020.09.10.292433

**Authors:** Kyrylo Pyrshev, Semen Yesylevskyy, Mikhail Bogdanov

**Author notes:** Authors contributed equally.

## Abstract

Identification, visualization and quantitation of cardiolipin (CL) in biological membranes is of great interest due to important structural and physiological roles of this lipid. Selective fluorescent detection of CL using non-covalently bound fluorophore TTAPE-Me (1,1,2,2-tetrakis[4-(2-trimethylammonioethoxy)-phenylethene) has been recently proposed. However, this dye was only tested on wild-type mitochondria or liposomes containing neglegible amounts of other anionic lipids, such as PG and PS. No clear preference of TTAPE-Me for binding to CL compared to PG and PS was found in our experiments. The shapes of the emission spectra for these anionic phospholipids were also found to be indistinguishable. Our experiments and complementary molecular dynamics simulations suggest that fluorescence intensity of TTAPE-Me is regulated by dynamic equilibrium between emitting dye, bound to anionic lipids by means of unspecific electrostatic attraction, and non-emitting dye aggregates in aqueous solution. Therefore, TTAPE-Me is not suitable for detection, visualization and localization of CL in the presence of PS and PG present in physiological amounts in the membranes of eukaryotic and prokaryotic cells, respectively.

## Introduction

Cardiolipin (CL) imparts cell membranes with a unique set of physical and chemical characteristics due to its unique chemical and structural properties: the high negative surface charge and small cross section of its head group relative to the cross section of four acyl chains [1]. The functions of CL are diverse. Besides crucial role in bioenergetics, including mitochondrial membrane morphology [2], CL is associated with many cellular and protein abnormalities and human diseases [3, 4]; involved in stabilization and activation of proteins in energy transducing membranes [5–8]; produces polar lipid microdomains [1, 9, 10]; serves as a targets for antimicrobials [11, 12]; controls distribution of bacterial cell division proteins [13]; sequester antibiotics apart of the division septum in bacteria [11, 12]; changes ordering and rigidity of the lipid bilayer [14]; contributes to bacterial adaptation to environmental stress [15, 16]; connects size and growth rate of bacterial cells [17] probably via its asymmetric distribution [18]; drives generation of outer membrane-derived vesicles by pathogenic bacteria [19]; serves as a signal for mitophagy [20, 21]; triggers production of antiphospholipid antibodies in pre-apoptotic cells due to its exposure to the cell surface [22, 23].

Despite the growing number of studies, the distribution of CL between the membrane monolayers in either bacterial or eukaryotic cells remains also unknown. Current methods of quantifying and visualizing CL *in vitro* and *in vivo* are hampered by the lack of specificity to this particular lipid. 10-N-nonyl acridine orange (NAO) is widely used for non-covalent fluorescent detection of CL in bacteria and mitochondria. NAO exhibits a characteristic red-shift of emission spectra, which was previously considered to be unique for the interaction between this probe and CL. However the specificity of NAO towards CL has been questioned recently. The characteristic red shift of NAO bound to PG and PS under their near-physiological concentration has been undoubtedly documented and the lack of specificity in both mitochondria [24], Gram-negative [10], Gram-positive bacteria [25] and archaea [26] was reported. Thus selective non-permeant vectorial (leaflet-specific) probes are of great demand for detection, localization and mapping an asymmetric distribution of CL.

Recently, a water soluble TTAPE-Me dye was proposed as a promising CL-specific fluorescent probe suitable for CL detection and sidedness studies [27, 28]. TTAPE-Me is non-fluorescent in aqueous solutions. Its fluorescence is turned on in CL-containing liposomes in concentration-dependent manner, while very weak fluorescence is observed in vesicles containing PC and PE lipids only [27].

Although TTAPE-Me is claimed to be selective to CL and is marketed in this way, this probe has never been tested properly for interactions with other anionic lipids. Indeed, in seminal work of Leung et al. only the wild-type mitochondria and liposomes representing a mitochondrial lipid composition were used [27]. CL is a predominant anionic lipid in mitochondria while other anionic lipids (PS, PI and PG) are found in very small percentage. Thus mitochondrial membranes and its lipid mimetics alone are not suitable for determining CL-specificity of the dye. Obviously these results are not relevant for the membranes of Gram-negative and Gram-positive bacteria, which are rich in PG, and mammalian plasma membranes where PS is abundant. Surprisingly, TTAPE-Me was never tested for equimolar concentrations of CL and other anionic lipids, particularly PG and PS.

It was reported recently that TTAPE-Me responds similarly to PG and CL [28], but these experiments were performed in the presence of drastic excess of the probe. As a result it is not known if TTAPE-Me is suitable for CL detection in PG-rich bacterial membranes and mammalian plasma membranes where PS is one of the most abundant lipids.

To fill this knowledge gap, we performed thorough investigation of TTAPE-Me combining fluorescence measurements with all-atom molecular dynamics (MD) simulations. We first assessed dye fluorescence in organic solvents with different degree of polarity and simulated its behavior in water. Than we measured dye fluorescence in liposomes constituted from major anionic lipids CL, PG and PS pristine or mixed with PE in different proportions. We also simulated interaction TTAPE-Me with lipid bilayers of corresponding compositions. This complementary approach allowed us (1) to elucidate the molecular mechanism of TTAPE-Me turn-on fluorescence in lipid membranes; (2) to show that fluorescence intensity of TTAPE-Me in different organic solvents is inconsistent with previously proposed emission mechanism [27]; (3) to demonstrate that this probe is not CL-specific and binds promiscuously to main anionic phospholipids.

## Results

Both the excitation and emission of TTAPE-Me were examined in different aqueous and organic solvents because an original report [27] considered excitation in water only.

The probe was essentially non-emissive in water, however the fluorescence increases drastically with a decrease of dielectric constant of the solvent (Fig 1). Although there are unsystematic deviations from this trend caused by the chemical nature of the solvent there is a clear indication that TTAPE-Me is more fluorescent in less polar environment. In this case of compounds of the same chemical nature (a series of monoatomic alcohols) there is a perfect linear dependence of the logarithm of emission intensity on dielectric constant (Fig. 1D).

**Figure 1.**
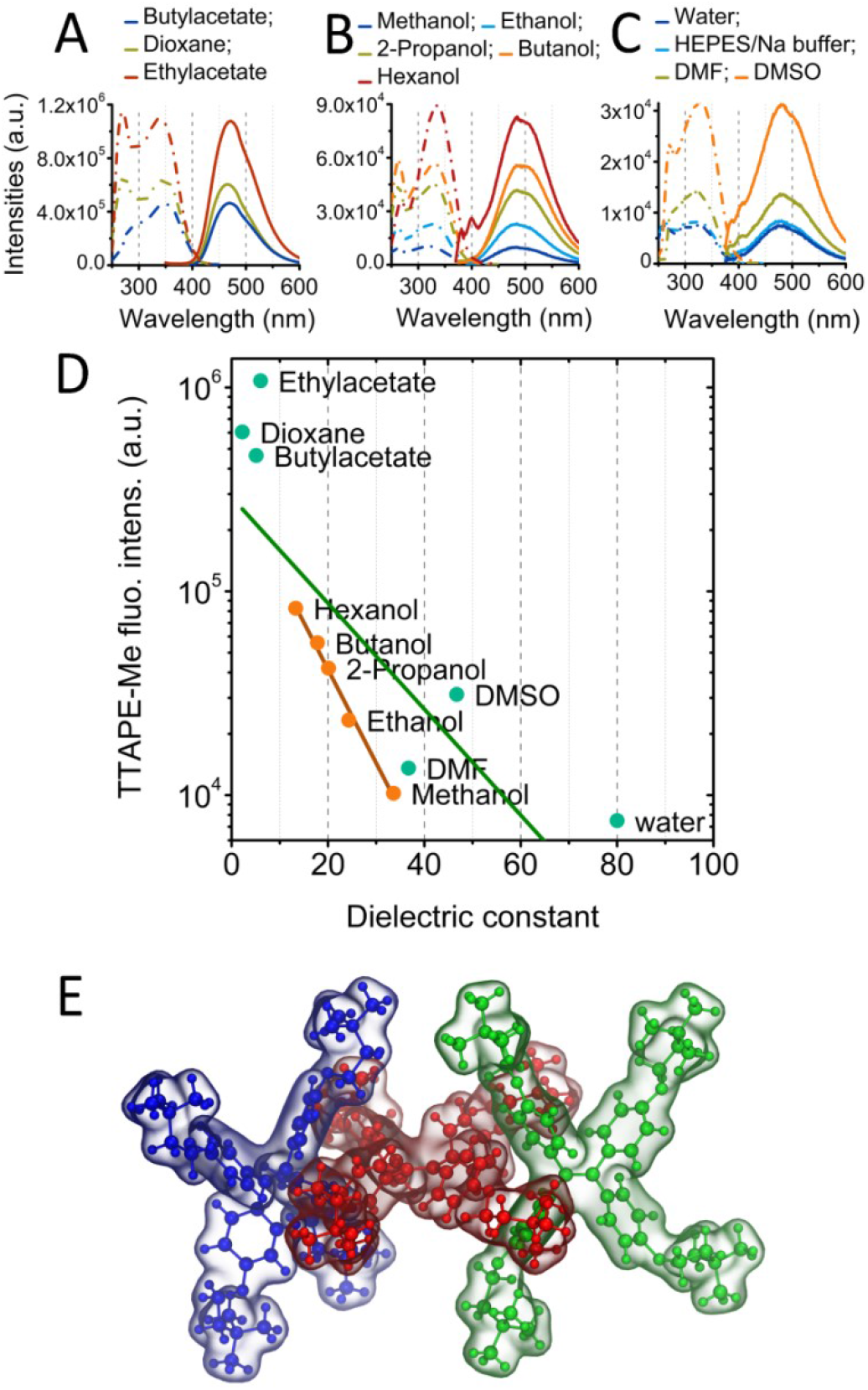
Fluorescent response of 20 μM TTAPE-Me probe to different solvents. (A,B,C) Excitation (dotted lines) and emission (solid lines) spectra in various solvents. (D) The maximal fluorescent intensities (in log scale) as a function of the dielectric constant of the solvent. Green line shows linear fit of all point. Brown line shows linear fit for monoatomic alcohols only. (E) Typical structure of the TTAPE-Me trimeric aggregate observed in MD simulations.

To understand this puzzling behavior of the dye we performed MD simulations of TTAPE-Me in aqueous solution. It appears that TTAPE-Me molecules in water are prone to fast spontaneous aggregation and formation of transient dimers and trimers. All aggregates adopt a “cogwheel topology” where the chromophores are positioned in a criss cross manner so that their hydrophobic central parts are in direct contact (Figure 1E). This 3D arrangement may minimizes the hydrophobic mismatch of the chromophores by hiding them from polar solvent. This finding allows us to hypothesize that very low fluorescent signal of TTAPE-Me in water is caused by the self-quenching in cogwheel-like aggregates. In more hydrophobic solvents the propensity of forming aggregates would be much lower due to smaller hydrophobic mismatch. As a result mostly monomeric probes would exist leading to intensive fluorescence.

Interestingly, the Leung et al. previously suggested the opposite mechanism where the probe is monomeric in water and the monomers are non-fluorescent [27]. However, our data disfavor this hypothesis.

In order to study interaction of TTAPE-Me with different anionic phospholipids we created SUVs consisting of CL, PG or PS alone or in binary mixture with PE at different mole ratios and measured the fluorescence emission intensities.

The optimal concentration of TTAPE-Me was determined by titration of different probe/lipid ratios (Figure 2C). The saturation of the signal starts at the ratio 20:1 for all lipids. Thus this ratio, corresponding to dye concentration of 10 μM, was used for all further experiments.

**Figure 2.**
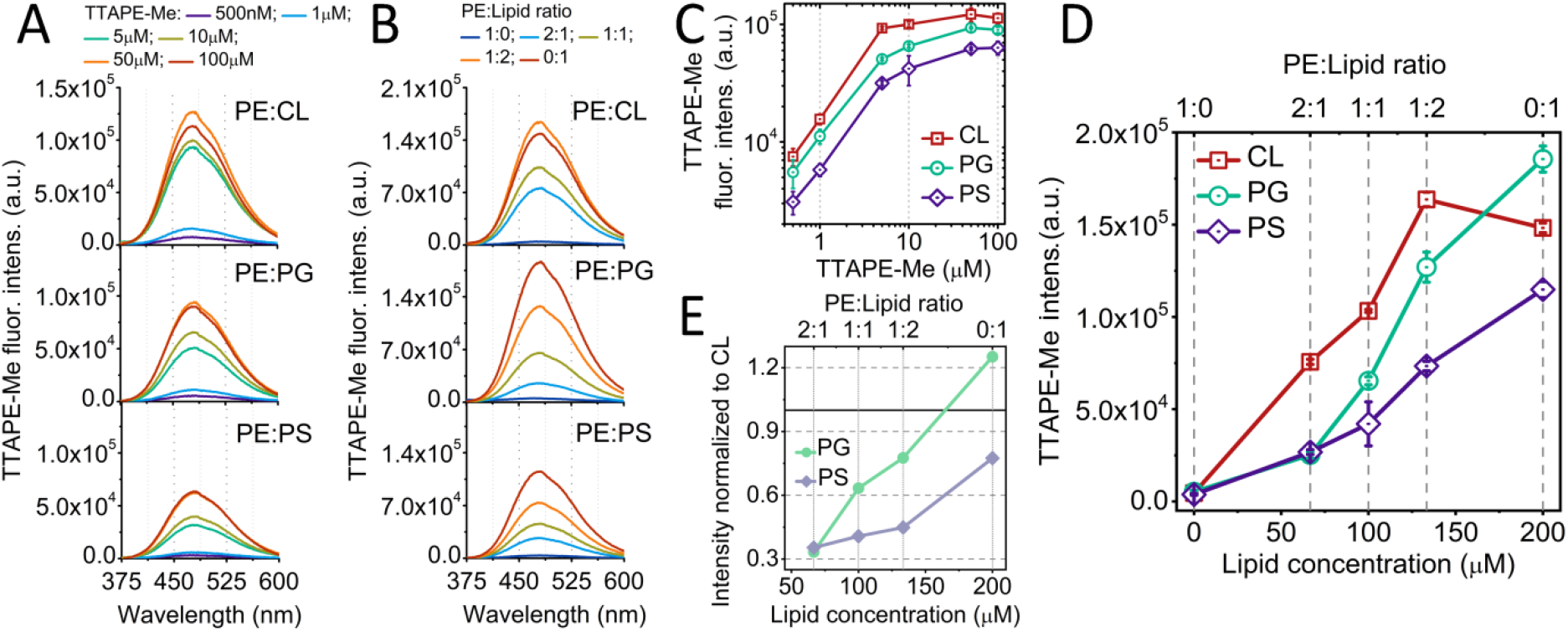
Fluorescent spectra of TTAPE-Me in 200 μM SUVs composed of PE mixed with CL, PG or PS in different proportions. (A) response of SUVs composed of PE:Lipid=1:1 mixtures (where “Lipid” refers to CL, PG or PS) to increasing concentration of TTAPE-Me. (B) response of SUVs with 10 μM of TTAPE-Me to increasing proportion of CL, PG and PS. (C) Fluorescence intensity as a function of TTAPE-Me concentration corresponding to A. (D) Fluorescence intensity as a function of concentrations of CL, PG and PS in model SUVs corresponding to B. Absolute concentration in shown on the bottom axis while the relative proportion of the lipid in relation to PE is shown on the top axis. (E) The ratios of fluorescent intensity in PG and PS in comparison to CL based on E. The axes notations are the same as on E. Data on C and D are represented as the mean±SD. Excitation at 325nm wavelength was used for all shown spectra.

Figure 2A and B show that the shapes of emission spectra for pure CL, PG and PS as well as for their binary mixtures with PE are indistinguishable. It is necessary to note that some published experiments [28] were performed in the presence of drastic excess of the probe and the background signal was not subtracted properly from the emission spectra. In such cases spurious differences in the shapes of spectra could be interpreted incorrectly as apparent selectivity of TTAPE-Me to particular lipids.

There is no significant fluorescence signal in pure PE which allows using mixtures with this aminolipid for measuring concentration dependencies for CL, PG and PS. The fluorescent intensities depend both on the nature of the anionic lipid and its content (Fig 2D). CL produces stronger response than PG and PS for concentrations up to 150 μM but the ratio of intensities never exceeds the factor of 3 (Fig 2E). For concentration of 200 μM the fluorescence of PG becomes stronger than CL (Fig 2D and E).

In order to rationalize observed results we performed MD simulations of monomeric TTAPE-Me with lipid bilayers made of CL, PG, PS and PC/PE. Analysis of trajectories shows that the monomeric probe does not interact with any of the studied lipids in specific manner. Only unspecific transient complexes of the probe with the lipids are formed.

The visual inspection (Fig. 3A,B) and the distance distributions **(Fig 3C)** show that the probe is mostly adsorbed on the membrane surface in CL and PG containing membranes, while it is less frequently bound to the PS membrane and only occasionally contacts the PC/PE membrane in the course of random diffusion. The probe molecule adsorbs on top of the lipid head groups. Multiple sorption/desorption events are observed in the course of simulations with one or more choline groups of TTAPE-Me being in contact with lipid head groups (Fig 3A,B). The probe does not penetrate into the bilayer and does not intercalate between the lipid head groups.

**Figure 3.**
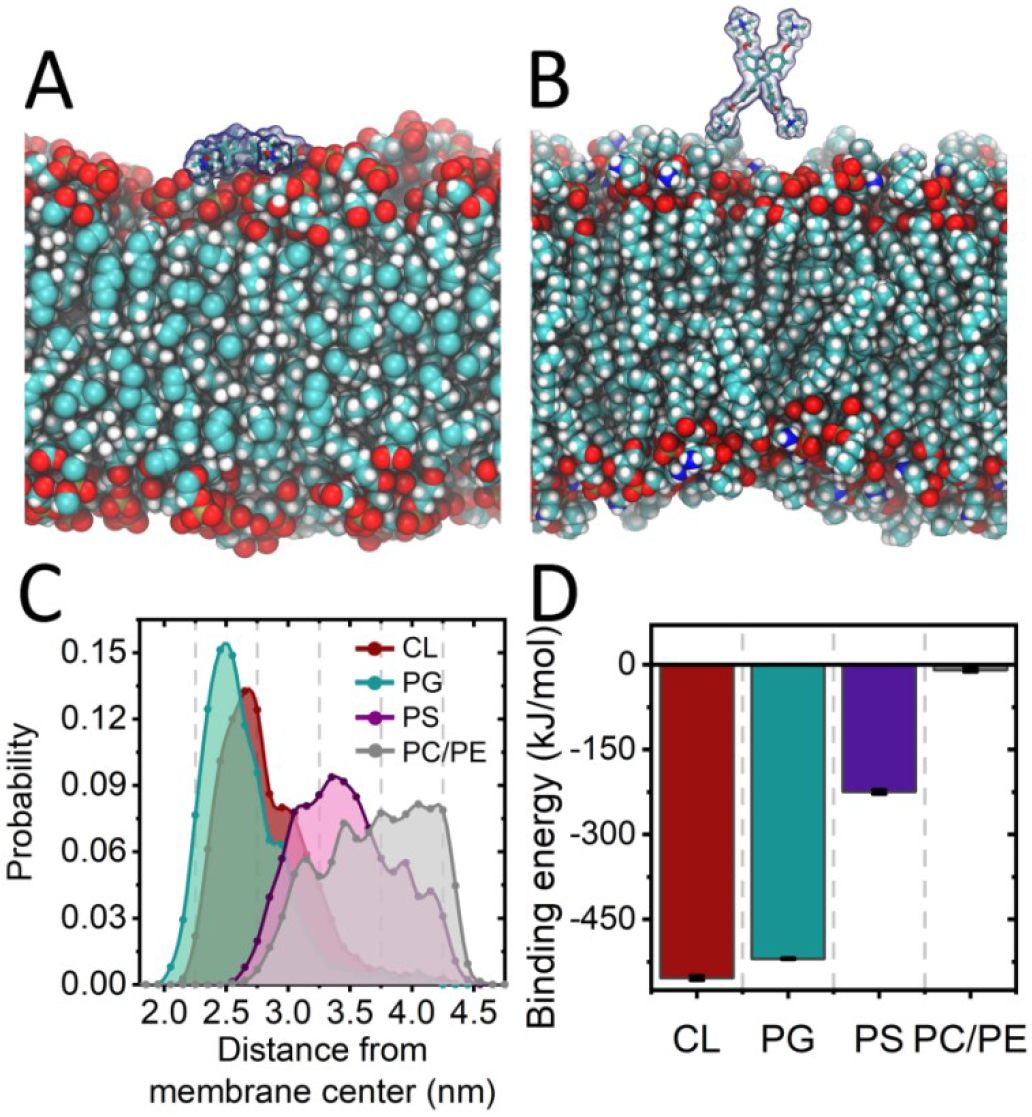
Typical snapshots of equilibrated MD trajectories for CL (A) and PC/PE (B) lipid bilayers. The lipids are shown in space-fill representation. The TTAPE-Me monomer is shown as balls and sticks with semi-transparent molecular surface. (D) Distributions of the distances from TTAPE-Me monomer to the center of lipid bilayer in MD simulations. (E) Interaction energies of TTAPE-Me monomer with different lipid bilayers computed over equilibrated parts of MD trajectories. Error bars correspond to the standard error.

Interaction energy between the probe and the membrane is dominated by electrostatic interactions and correlates strongly with lipid net charge and the charge of the head group (Fig 3D, Table 1). The probe is attracted to the membrane by the lipid net charge while the strength of binding is likely to be further tuned by the part of the head group located above the phosphate moiety, which is in direct contact with the probe. Indeed, the probe binds strongly to CL and PG which lack any charged moieties above the phosphate. The binding to PS, which possesses a serine group, is weaker due to unfavorable interactions with exposed primary amine. Finally, there is no interaction at all with PC and PE because these lipids hold net neutral charge and own an exposed amine (choline and ethanolamine, respectively), which repel strongly cationic TTAPE-Me molecule.

**Table 1.**
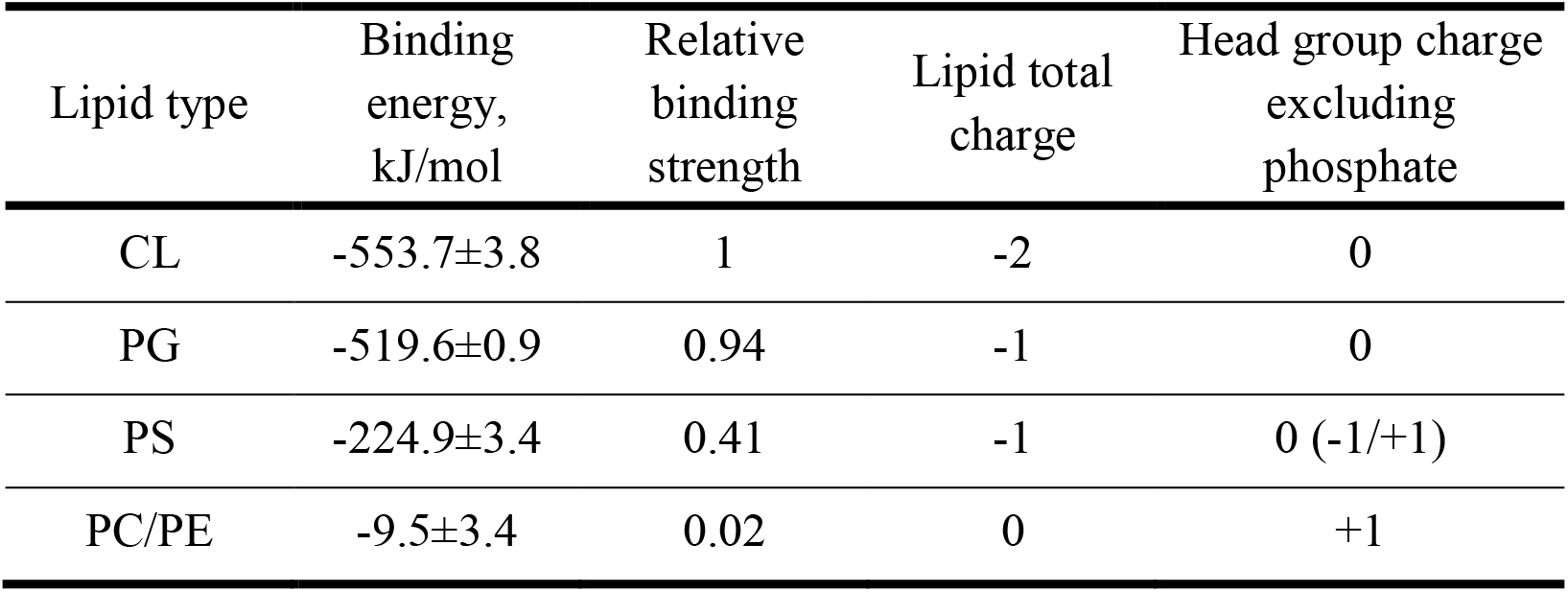
Binding of TTAPE-Me to the lipid bilayer as a function of the lipid total charge and the charge of head group above the phosphate moiety. The mean binding energies and the standard errors are the same as in Fig. 3D. Relative binding strength is computed in comparison to CL.

| Lipid type | Binding energy, kJ/mol | Relative binding strength | Lipid total charge | Head group charge excluding phosphate |
| --- | --- | --- | --- | --- |
| CL | -553.7±3.8 | 1 | -2 | 0 |
| PG | -519.6±0.9 | 0.94 | -1 | 0 |
| PS | -224.9±3.4 | 0.41 | -1 | 0 (-1/+1) |
| PC/PE | -9.5±3.4 | 0.02 | 0 | +1 |

The interaction energies of monomeric probe with different lipids, obtained in MD simulations, are in perfect agreement with observed fluorescence intensities in the corresponding SUVs, which decrease in the same manner CL ≥ PG > PS > PE. This suggests convincingly that the fluorescence in the presence of anionic lipids is facilitated by TTAPE-Me monomers, rather than aggregates as was initially hypothesized by Leung et al. [27].

Our data allows formulating the following hypothesis of TTAPE-Me action in the presence of anionic lipid membranes. The fluorescence comes from the probe monomers unspecifically adsorbed on top of the membrane surface by means of unspecific electrostatic attraction modified slightly by the nature of exposed head groups. There is a dynamic equilibrium between emitting membrane-adsorbed monomers and non-emitting self-quenched aggregates in the bulk solvent. The more probe molecules adsorb on the membrane surface, the less aggregates remain in solution and the stronger is the fluorescence signal. This scheme describes all our experimental and simulation data in a simple and elegant way and explains observed fluorescent intensities in a row CL ≥ PG > PS > PC/PE.

## Conclusions

Our data show that TTAPE-Me is not selective to CL or to any other anionic lipid. This probe is sensitive to the net change of the membrane surface and the charge of exposed moieties of the lipid head groups in generic manner. The most plausible mechanism of this sensitivity is dynamic equilibrium between self-quenching probe aggregates in solution and the fluorescent monomers adsorbed on the lipid head groups by means of unspecific electrostatic interactions. The fluorescent responses of TTAPE-Me for different anionic lipids are comparable and do not allow discriminating between CL, PG and PS lipids by either shape of the spectra or emission intensities.

TTAPE-Me could still be relevant for detecting CL in in mitochondria, which lack significant amounts of any other anionic lipids. However, this probe is expected to bind promiscuously to all anionic phospholipids in eukaryotic cells, Gram-positive and Gram-negative bacteria and the cell-derived vesicles.

Marketing of TTAPE-Me as “Cardiolipin probe” is highly misleading because this probe is not suitable for selective labeling of CL in the membranes where other anionic lipids are present in non-negligible physiological amounts.

These results should be taken into consideration when interpreting past and future results of CL detection and localization studies with TTAPE-Me probe *in vivo* and *in vitro*.

## Materials and methods

### Molecular dynamics simulations

The topology of TTAPE-Me probe was generated with CHARMM GUI ligand builder [29, 30]. In order to study behavior of the probe in water ten TTAPE-Me molecules were put randomly into 5×5×5 nm cubic box and solvated with ~3500 water molecules. The 11 Na^+^ and 51 Cl^−^ were added to neutralize the net positive charge of the probe and to achieve total ionic concentration of 0.15 M.

In order to simulate interactions of TTAPE-Me with lipid membranes five lipid bilayers were constructed using CHARMM GUI membrane builder [29, 30] with the composition detailed in Table 2. Pure PE does form SUVs in experiments but normally does not exist as a stable flat bilayer thus a mixed PC/PE system was used in simulations.

**Table 2.** Composition of the lipid bilayers used. Lipid names correspond to the residues names in CHARMM36 force field [31]. TOCL2 is 1′,3′-bis[1,2-dioleoyl-sn-glycerol-3-phospho-]-sn-glycerol; DOPC is 1,2-dioleoyl-sn-glycero-3-phosphocholine, DOPE is 1,2-dioleoyl-sn-glycero-3-phosphoethanolamine; DOPG is 1,2-dioleoyl-sn-glycero-3-phospho- (1’-rac-glycerol); DOPS is 1,2-dioleoyl-sn-glycero-3-phospho-L-serine.

| System name | Lipids | Ions | Water |
| --- | --- | --- | --- |
| CL | TOCL2: 64 | Na<sup>+</sup>: 142<br>Cl<sup>-</sup>: 18 | 6396 |
| PC/PE | DOPC: 64<br>DOPE: 64 | Na<sup>+</sup>: 13<br>Cl<sup>-</sup>: 17 | 6396 |
| PG | DOPG: 128 | Na<sup>+</sup>: 141<br>Cl<sup>-</sup>: 17 | 6396 |
| PS | DOPS: 128 | Na<sup>+</sup>: 142<br>Cl<sup>-</sup>: 18 | 6396 |

All bilayers were built with 64 lipids per monolayer, except CL which had 32 lipids per monolayer due to double size of each lipid molecule. Bilayers were solvated with 50 water molecules per lipids. The number of ions corresponds to 0.15 M of NaCl with additional counter ions added to neutralize the system.

MD simulations were performed in Gromacs [32] version 2019.2 in NPT ensemble at the pressure of 1 atm maintained by Parrinello-Rahman barostat [33]. Isotropic pressure coupling was used for the probe in water and the semi-isotropic coupling for the membrane simulations. The Verlet cutoff scheme was used [34]. Force-switch cut-off of the Van der Waals interactions was used in the region between 1.0 and 1.2 nm. Long range electrostatics was computed with the PME method [35] with the cut-off of explicit short-range electrostatic interactions at 1.2 nm. Velocity rescale thermostat [36] was used at the temperature of 300K.

The probe in water was simulated for 100 ns. All bilayers were pre-equilibrated for 100 ns. After that the probe was inserted at the distance of ~4 nm from the membrane center and the system was simulated for another 100 ns. The interaction energy between the probe and the membrane was monitored during the whole production run.

CHARMM36 force field [31] was used for all components of the system. An integration step of 2 fs was used in all simulations with the bonds to hydrogen atoms converted to rigid constraints. Analysis was performed by the custom scripts based on Pteros molecular modeling library [37]. VMD 1.9.3 was used for visualization [38].

### Reagents

TTAPE-Me (Biovision); phosphatidylethanolamine total extract from E. coli (Avanti); cardiolipin total extract from E.coli (Avanti); phosphatidylglycerol total extract from E.coli (Avanti); phosphatidyserine extract from porcine brain (Avanti).

### SUV preparation

The small unilamellar vesicles (SUVs) were prepared by sonication method as it was described elsewhere [39, 40]. Briefly, the stock solution of lipids was dried in sample tube till they form lipid film and then diluted in 10 mM Na/HEPES buffer (pH 7.4) to obtain 1 mM concentration. Formation of the aggregates or multilamellar vesicles results in white color suspension. Further sonication with the probe tip resulted in a transparent opalescent solution of small unilamellar vesicles. The size of SUVs was measured by laser correlation spectroscopy (Malvern) and it corresponds to 40±7nm. The SUVs composed of PE contained more aggregates but due to the overall low response of TTAPE-Me to this lipid, this was not considered significant for the current study.

### Labelling procedures

Initially, the stock powder of TTAPE-Me probe was diluted in DMSO. For monitoring of TTAPE-Me response to different solvents, the probe in DMSO was directly added to the respective solvent to receive 20 μM final concentration of the probe and >0.25% of DMSO (except the samples with pure DMSO).

To label the SUVs, TTAPE-Me was diluted in 600 μL 10mM HEPES/Na buffer (pH 7.4) and added to 150μL of 1mM mixture of the lipids. As the result, 200μM solution of the lipids (>0.25% of DMSO) was received. The mixture was incubated during 10 min on a dark at room temperature before the measurements.

### Spectroscopy

Spectra were recorded in half-micro cuvette with PTI Quantamaster 40 spectrofluorometer at 25°C. 3 nm slits on excitation and emission were applied to receive reasonably high signal.

To minimise the influence of autofluorescence and scattering on the probe response, the spectra of the TTAPE-ME in SUVs or solution were normalized on the curve provided by the respective vesicles or solvent without the probe.

## Contribution

MB initiated the research. MB, SY and KP designed the experiments. KP performed *in vitro* experiments. SY performed *in silico* experiments. All authors interpreted the data and wrote the manuscript. The authors declare no conflict of interest.

## Acknowledgement

This work was supported by NATO Science for Peace and Security Programme-SPS 985291 (to MB), the European Union’s Horizon 2020 research and innovation programme under the Marie Skłodowska-Curie grant agreements No. 690853 (to MB and SY), Program of Competitive Growth of Kazan Federal University (to MB); KP and SY were supported also by stipend from NATO Science for Peace and Security Programme-SPS 985291. SY was supported by the European Union’s Horizon 2020 research and innovation programme under the Marie Skłodowska-Curie grant agreement No. 796245.

## Notes

### Competing Interest Statement

The authors have declared no competing interest.

